# Cocaine-induced neuron subtype mitochondrial dynamics through Egr3 transcriptional regulation

**DOI:** 10.1101/2020.06.27.175349

**Authors:** Shannon Cole, Ramesh Chandra, Maya Harris, Ishan Patel, Torrance Wang, Hyunjae Kim, Leah Jensen, Scott J Russo, Gustavo Turecki, Amy M Gancarz-Kausch, David M Dietz, Mary Kay Lobo

## Abstract

Mitochondrial function is required for brain energy homeostasis and neuroadaptation. Recent studies demonstrate that cocaine affects mitochondrial dynamics and morphological characteristics within the nucleus accumbens (NAc). Further, mitochondria are differentially regulated by cocaine in dopamine receptor-1 containing medium spiny neurons (D1-MSNs) vs dopamine receptor-2 (D2)-MSNs. However, there is little understanding into cocaine-induced transcriptional mechanisms and their role in regulating mitochondrial processes. Here, we demonstrate that cocaine enhances binding of the transcription factor, early growth response factor 3 (Egr3), to nuclear genes involved in mitochondrial function and dynamics. Moreover, cocaine exposure regulates mRNA of these mitochondria-associated nuclear genes in both contingent or noncontingent cocaine administration and in both rodent models and human postmortem tissue. Interestingly, several mitochondrial nuclear genes showed distinct profiles of expression in D1-MSNs vs D2-MSNs, with cocaine exposure generally increasing mitochondrial-associated nuclear gene expression in D1-MSNs vs suppression in D2-MSNs. We further show that blunting Egr3 expression in D1-MSNs blocks cocaine-enhancement of the mitochondrial-associated transcriptional coactivator, peroxisome proliferator-activated receptor gamma coactivator (PGC1α), and the mitochondrial fission molecule, dynamin related protein 1 (Drp1). Finally, reduction of D1-MSN Egr3 expression attenuates cocaine-induced enhancement of small-sized mitochondria, causally demonstrating that Egr3 regulates mitochondrial morphological adaptations. Collectively, these studies demonstrate cocaine exposure impacts mitochondrial dynamics and morphology by Egr3 transcriptional regulation of mitochondria-related nuclear gene transcripts; indicating roles for these molecular mechanisms in neuronal function and plasticity occurring with cocaine exposure.

## Introduction

Mitochondria are chiefly responsible for energy production in cells; a basic requirement for mammalian life and cellular adaptation. Recent studies revealed the important relationship of mitochondrial programming, machinery, and adaptations in brain responses to drugs of abuse in humans and in rodent models (1–3). In the nucleus accumbens (NAc), mitochondrial fission or molecules related to mitochondrial dynamics and function are altered by cocaine (1,2).

The NAc is a critical hub involved in motivation for various rewards, such as food and drugs of abuse, as well as fear and aversion (4,5). The NAc is primarily comprised of medium spiny neurons (MSNs) of two distinct subpopulations: D1-MSNs and D2-MSNs (6,7). D1 and D2-MSN activity and morphology are altered across multiple pathological states, such as addiction, stress, depression, and pain (8–12).

NAc neurobiological molecular mechanisms involved in neuroplasticity often differ between D1- and D2-MSNs in response to cocaine (12–20). Further, patterns of mitochondria-related gene regulation can be distinct and opposing between these neuron populations (1,2). For instance, following cocaine exposure or self-administration the mitochondrial fission protein, dynamin related protein (Drp1), and a transcriptional coactivator molecule involved with mitochondrial dynamics, peroxisome proliferator-activated receptor gamma coactivator (PGC1α), are enhanced in D1-MSNs and decreased in D2-MSNs (1,2,21,22). Interestingly, recent studies have discovered that mitochondrial fission via Drp1 is required for neuronal spine formation (24,24); a key morphological adaptation for enhancing synaptic strength that may underlie craving and seeking of drugs after consumption has ceased (25–33).

The NAc contains other drug-responsive molecular mechanisms that may act upstream of mitochondrial related molecules. One such transcription factor is early growth response factor 3 (Egr3), which regulates several cocaine-associated genes, such as CREB, CamKIIα, and ΔFosB (14). Egr3 displays opposing function in D1-MSNs vs D2-MSNs: Egr3 is upregulated in D1-MSNs following repeated cocaine and is necessary for cocaine conditioned place preference (CPP) and cocaine induced psychomotor locomotion. Conversely, D2-MSNs have reduced Egr3 following repeated cocaine and enhancing D2-MSN Egr3 causes reduced cocaine CPP and psychomotor locomotion. Effects of Egr3 in D2-MSNs are also observed after long term abstinence, but in a sex-specific manner (34). Further, Egr3 action on cocaine-regulated behaviors could occur through mitochondrial processes, since cocaine exposure enhances Egr3 binding to the PGC1α promoter in NAc (2). Collectively, these findings suggest an important link between Egr3 transcription, molecules involved with mitochondrial processes, and facilitation of cocaine-induced behaviors.

Relatively little is known about how drug exposure alters mitochondrial morphological characteristics by transcriptional regulation of mitochondria-related genes. This study assayed the impact of cocaine exposure on NAc Egr3 regulation of multiple downstream target nuclear genes involved with mitochondrial-related processes and mitochondrial morphology. Mitochondria-related nuclear genes were assayed following contingent or non-contingent cocaine exposure in rodents and were also examined in postmortem NAc of cocaine dependents. Further, mitochondrial-related nuclear genes were examined in NAc MSN subtypes after exposure to cocaine and when Egr3 is genetically disrupted. Mitochondrial-related nuclear genes include: the transcription factors nuclear respiratory factor 1 and 2 (Nrf1 and Nrf2); a transcriptional coactivator of mitochondrial related nuclear genes, PGC1α; mitochondria-specific transcription factors a and b1 (Tfam and Tfb1); the mitochondrial trafficking membrane protein, translocase of the outer mitochondrial membrane 20 (Tomm20); the mitochondrial fission regulator, Drp1; and a mitochondrial DNA polymerase subunit gamma, Polγ (a.k.a. POLG). Together, these studies examine the impact of cocaine exposure on mitochondrial dynamics and morphology through Egr3 transcriptional regulation of mitochondria-related nuclear gene transcripts.

## Materials and Methods

### Animals and Human Postmortem Samples

All animal experiments and handling followed guidelines set by the Institutional Animal Care and Use Committee (IACUC) at The University of Maryland School of Medicine and The University at Buffalo, The State University of New York. In both institutions, the health status of test animals was closely monitored by onsite veterinary staff and the experimenters throughout all experiments. Mice were maintained on 12h light/dark cycle with *ad libitum* food and water. All mice were group housed (3-5 per cage).

For chromatin immunoprecipitation (ChIP) experiments male C57BL/6J mice were obtained from Jackson Laboratory for cocaine treatment (n=35) and saline control groups (n=30) and were pooled with 5 mice per sample (n=6 saline control samples, n=7 cocaine treatment samples).

C57BL/6J male mice (n=6 for saline control, n=7 for cocaine treatment) used in quantitative real time polymerase chain reaction (qRT-PCR) quantification of whole NAc tissue following i.p. cocaine were obtained from Jackson Laboratory.

This study utilized both male and female D1-Cre hemizygote or D2-Cre hemizygote bacterial artificial chromosome transgenic mice from GENSAT (D1-Cre hemizygote line FK150; D2-Cre hemizygote line ER44)(35) (www.gensat.org) on a C57BL/6J background.

Male and female homozygous RiboTag (RT) mice on a C57BL/6J background, expressing a Cre-inducible HA-Rpl22 (Sanz et al., 2009) were crossed to D1-Cre or D2-Cre hemizygote mouse lines to generate D1-Cre-RT and D2-Cre-RT male and female mice (1), and used for cell subtype–specific ribosome-associated mRNA isolation. Tissue from RiboTag experiments came from mice also published elsewhere (36). RiboTag testing comprised of samples from D1-Cre-RT (saline control: n=16 females, n=8 males; cocaine treatment: n=8 females, n=16 males) and D2-Cre-RT mice (saline control: n=8 females, n=16 males; cocaine treatment: n=12 females, n=12 males), ages 8-10 weeks with 4 mice pooled per sample (D1-Cre-RT saline pooled samples: n=4 females, n=2 males; D1-Cre-RT cocaine pooled samples: n=2 females, n=4 males; D2-Cre-RT saline pooled samples: n= 2 female, n=4 female; D2-Cre-RT cocaine pooled samples: n=3 female, n=3 males). Animal numbers were determined based on previous studies (1,2,14).

D1-Cre-RT mice were used for qRT-PCR analysis for Egr3-miR experiments: SS-miR saline control (n=12 male, n=8 female), SS-miR cocaine treatment (n=12 male, n=12 female), Egr3-miR saline treatment (n=16 male, n=8 female), and Egr3-miR cocaine treatment (n=16 male, n=12 female). 4 mice were pooled per sample: SS-miR saline control (n=3 male, n=2 female), SS-miR cocaine treatment (n=3 male, n=3 female), Egr3-miR saline treatment (n=4 male, n=2 female), and Egr3-miR cocaine treatment (n=4 male, n=3 female).

For morphological characterization of mitochondria after Egr3-miR or SS-miR control, male and female D1-Cre mice age 10-12 weeks were used (SS-miR saline: n=4 male; SS-miR cocaine: n=4 male; Egr3-miR saline: n=2 female, n=3 male; Egr3-miR cocaine: n= 2 female, and 3 male).

Naive male Sprague-Dawley rats (250-275g at the beginning of the experiment), used for self-administration cDNA, were maintained on a 12h reverse light/dark cycle ad libitum food and water. After catheterization surgery, rats were single housed for the duration of the experiments. For all *in vivo* studies, animals were randomly assigned to the treatment (n=6) or control group (n=5).

Human postmortem tissue was also used in (1) and was handled according to the same storage and treatment protocols. The cohort of cocaine-dependent individuals was comprised of 24 males and 2 females ages 20-53 years (see Additional File S1 for subject details). All subjects died suddenly without a prolonged agonal state or protracted medical illness. The causes of death were ascertained by the Quebec Coroner Office. A toxicological screen was conducted with tissue samples to obtain information on medication and illicit substance use at the time of death. Cocaine-dependent subjects included 14 individuals who met the Structured Clinical Interview for Diagnostic and Statistical Manual of Mental Disorders-IV (DSM-IV) Axis I Disorders: Clinician Version (SCID-I) criteria for cocaine dependence. The control group included 12 subjects with no history of cocaine dependence or major psychiatric diagnoses. The processing of the tissue is outlined in previous publications (37,38). Hemispheres were immediately separated by a sagittal cut into 1cm-thick slices and placed in a mixture of dry ice and isopentane (1:1; vol:vol) and then stored at −80°C. Tissue weighing approximately 150 mg was punched from the NAc, homogenized in 5 ml of ddH2O (pH adjusted to 7.00) and centrifuged for 3 min at 8,000g and 4°C. The pH of the supernatant was measured in duplicate (Thermo-Electron Corporation). RIN determination was performed by isolating total RNA using TRIzol (Invitrogen) followed by analysis with an Agilent 2100 Bioanalyzer.

### Mouse stereotaxic surgery

Mice were anesthetized using 4% isoflurane in a small induction chamber. After initial induction, isoflurane was maintained at 1-2% for the remainder of the surgery. Animals were placed in a stereotaxic frame and their skull was exposed. 33-gauge Hamilton syringe needles were used to inject Cre-inducible double inverted open (DIO) reading frame adeno-associated viruses (AAV) (1) into the NAc of test animals. To label mitochondria, AAV-DIO-mito-dsRed was injected in tandem with either AAV-DIO-Egr3-miR-IRES-mCitrine for Egr3 knockdown or AAV-DIO-Scramble(SS)-miR-IRES-mCitrine for viral control in D1-Cre-RT mice for RiboTag experiments and D1-Cre mice for morphology experiments (1,2,14,39).

All experiments involving viral infusion had virus injected bilaterally into the NAc (anterior/posterior, AP+1.6; medial/lateral, ML ± 1.5; dorsal/ventral, DV-4.4, 10 degree angle). Mice were then returned to the vivarium for 2 weeks to allow for recovery and virus expression. All viruses were packaged in lab facilities as previously described (1).

### Cocaine injection and cocaine self-administration

C57Bl/6J, D1-Cre, D1-Cre-RT, and D2-Cre-RT mice received 7 daily intraperitoneal injections (i.p. of cocaine, 20mg/kg) or saline control injections in the home cage. NAc tissue for all experiments with mice was collected 24h after the last injection. Cocaine hydrochloride (Sigma) was dissolved in sterile saline, and the dose of cocaine was determined based on effects from previous studies (1,2,12–14,40).

Protocols for rat cocaine self-administration experiments were adapted from previously published in-house experiments (1,2,14,33,40). Shortly, 1 week following catheterization, rats went through 10 daily 2hr sessions of intravenous cocaine or saline self-administration (1mg/kg/infusion in sterile saline solution) at an FR1 schedule reinforcement and a 30s time out period between infusions. 24hrs after the last cocaine infusion, rat NAc tissue was collected.

### RiboTag and qRT-PCR procedure

NAc samples were pooled from four D1-Cre-RT or D2-Cre-RT male and female mice and RNA isolation from immunoprecipitated polyribosomes using primary mouse anti-HA antibody (Covance Research Products, Cat#MMS101R) and secondary antibody coated magnetic Dynabeads (Dynabeads protein G, Invitrogen) to pulldown the MSN specific RNA. Immunoprecipitated polyribosomes and non-immunoprecipitated input was prepared according to our previous studies (1,14,39). In brief, four 14-gauge NAc punches per animal (four animals pooled per sample) were collected and homogenized by douncing in homogenization buffer and 800 mL of the supernatant was added directly to the HA-coupled beads (Invitrogen: 100.03D; Covance: MMS-101R) for constant rotation overnight at 4°C. RNA from polyribosome immunoprecipitated samples or input was subsequently extracted using the Total RNA Kit (Omega) according to manufacturer’s instructions. RNA was quantified with a NanoDrop (Thermo Scientific).

For cDNA synthesis, qRT-PCR, and analysis we follow the steps as described previously (Chandra, Engeln et al., 2017; Chandra et al., 2015; See Additional File S2 for list of primers). Mouse and rat NAc tissue punches were collected 24 h after the last cocaine administration and stored at −80°C. For both rodent and human postmortem NAc tissue, total RNA was isolated by using Trizol (Invitrogen) homogenization and chloroform layer separation (37). A clear RNA layer was then processed (RNAeasy MicroKit, Qiagen) and analyzed with NanoDrop (Thermo Scientific). Next, 500ng of RNA was reverse transcribed to cDNA (qScript Kit, Bio-Rad). cDNA was diluted to 200μl, and 2μl was used for each reaction. Human mRNA integrity was determined on an Agilent Bioanalyzer. mRNA expression changes were measured using qPCR with PerfeCTa SYBR Green FastMix (Quanta). Quantification of mRNA changes was performed using the -ΔΔ C_T_ method, using glyceraldehyde 3-phosphate dehydrogenase (GAPDH) as a housekeeping gene. -ΔΔ C_T_ cDNA values that showed no detection were removed for specific genes. In qRT-PCR analysis, samples 2 standard deviations above or below the mean were excluded. Samples were also excluded with ΔΔ C_T_ values more than 30 or in the event that no ΔΔ C_T_ values were present after analysis.

### Chromatin immunoprecipitation

Fresh NAc punches were prepared for ChIP as previously described with minor modifications (2,14). Briefly, 4, 14-gauge NAc punches per C57BL/6J mouse (four animals pooled per sample) were collected, cross-linked with 1% formaldehyde and quenched with 2M glycine before freezing at −80°C. Before sample sonication, IgG magnetic beads (Invitrogen; sheep anti-rabbit, cat.# 11202D) were incubated with an anti-Egr3 (rabbit polyclonal, Santa Cruz Biotech, cat.# SC-191, 8–10μg per reaction) antibody overnight at 4°C under constant rotation in phosphate buffered saline (PBS)-based blocking solution (0.5% w/v BSA in 1X PBS). The chromatin was fragmented by sonication to an average length of 500-700bp. After sonication, equal concentrations of chromatin were transferred to new tubes and 100μl of the final product was saved for input controls. Following the washing and resuspension of the antibody–bead conjugates, antibody–bead mixtures were added to each chromatin sample (600μl) and incubated for ~16h under constant rotation at 4°C. Samples were then washed and reverse cross-linked at 65°C overnight and DNA was purified using a PCR purification kit (QIAGEN). After DNA purification, samples were used for qPCR analysis and normalized to their appropriate input controls as previously described(14,41,42). Rabbit IgG immunoprecipitation control sample was prepared by adding blocking buffer to magnetic beads and anti-Egr3 for appropriate enrichment of Egr3 ChIPs, >1.5 fold.

### Immunohistochemistry

Mice were perfused with 0.1M PBS followed by 4% paraformaldehyde (PFA). Brains were immersed in PBS-azide overnight. Brains were dissected with a vibratome (Leica) at 100 mm into PBS-azide for mitochondria.- Brain sections were blocked in 3% normal donkey serum with 0.3% Triton-X for 30 min at room temperature. Sections were then incubated overnight at 4°C in primary antibody (1:1500 chicken anti-GFP; Aves Labs, Cat#GFP-1020) diluted in 3% NDS and 0.3% tween 20 solution. On the second day, tissue sections were rinsed in PBS three times for 30 min per rinse followed by rinsing with PBS every hour at room temperature for 8 hours, then incubated overnight at 4°C in secondary antibody, 1:500 donkey anti-chicken-Alexa488 (Jackson ImmunoResearch cat# 703-545-155) diluted in PBS. After secondary incubation, sections were rinsed in PBS every hour for 8 hours. Slices were mounted, dried overnight in the dark, and cover slipped with mounting media (Aqua-Polymount; Polysciences, Inc.). DsRed was not amplified via immunostaining, as endogenous virus expression was sufficient for imaging.

### Mitochondrial Imaging and Analysis

Sections were imaged on a Leica SP8 confocal microscope. A total of 4-5 neurons were imaged per mouse. High-resolution Z-stacks images were obtained with 0.4μm increments using a 60x oil immersion objective with 2x digital zoom. Sections were scanned for neuron distal dendrites (over 100mm radius from the soma) and distal secondary dendrites (the first nearest branch from distal dendrite). Dendrites and mitochondria were 3-dimentionally reconstructed using Imaris 8.2 software (Bitplane), and multiple morphological parameters were analyzed, including mitochondrial length, density, index, and volume.

3D reconstructions of images were generated of each filter: red (mitochondria) and green (soma and dendrites). In short, mitochondria length was measured by using the BoundingBoxOO Length C setting that measures the length of the longest principal axis of the mitochondria. Dendritic length was determined by ‘measurement point’ tool in line pair mode. Mitochondrial volume was measured by quantifying total mitochondrial volume divided by total dendrite volume. Mitochondrial density was measured by quantifying the total number of mitochondria per 10μm length of dendrites. Dendritic mitochondrial indices were measured by quantifying the average mitochondrial length per 10μmlength of dendrite. Analysts were blinded to test conditions.

### Statistical analysis

GraphPad Prism 6.0 software was used for statistical analysis. Two-way ANOVA was used for mitochondrial size frequency analysis followed by Bonferroni post-hoc correction. Student’s t-test was used for all other experiments. Comparisons were tested for normalcy. Significance was established when p-values were below 0.05. All graphs represent mean ± standard error (SEM). For qRT-PCR data, Grubbs outlier test was performed on data with obvious outliers and no more than one animal or sample was removed per group. No statistically significant sex differences were observed in any applicable experiment and so all groups with both male and females were combined.

## Results

Egr3 binding sites are found on promoters of many nuclear, mitochondrial-related genes (gene-regulation.com, Alibaba). To determine if cocaine exposure alters Egr3 binding to the promoters of these mitochondrial-related nuclear genes we performed ChIP (Fig 1a). Following 7-days of i.p. cocaine injections (20mg/kg) NAc tissue was collected 24 hours after the last injection (Fig 1b), a time point shown to induce transcriptional and cellular plasticity in NAc (2,14,20,41,43,44). In bulk NAc tissue, enhanced binding of Egr3 was observed on the promoters of the mitochondrial fission molecule, Drp1, the transcription factor, nuclear respiratory factor, Nrf2, and the DNA polymerase gamma subunit, Polγ. This is in line with a previous study demonstrating enhanced Egr3 binding at the PGC1α promoter in NAc after repeated cocaine (2). These findings implicate cocaine-mediated Egr3 transcriptional regulation of mitochondrial-related nuclear genes.

**Figure 1.**
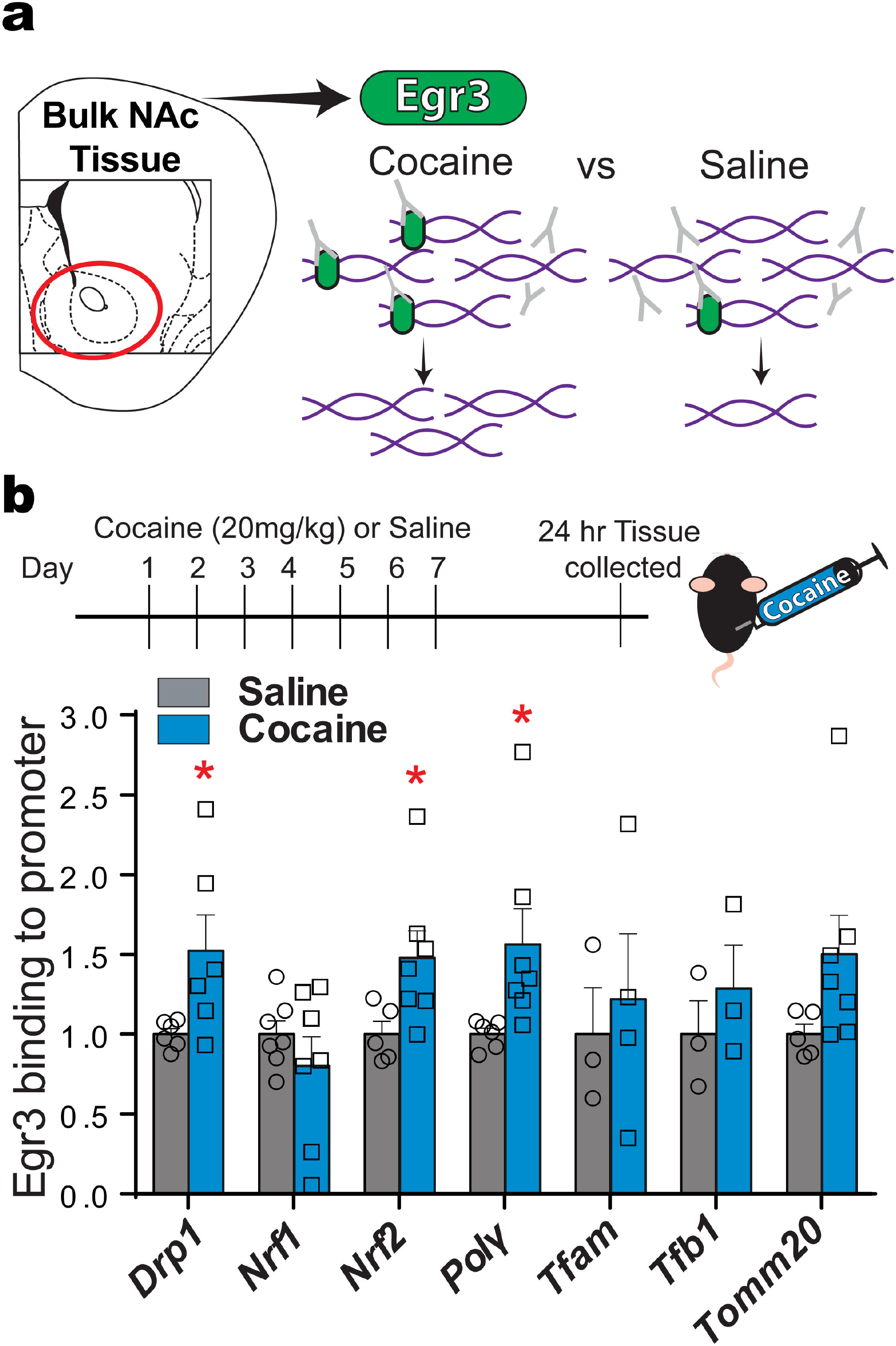
Cocaine-induced Egr3 binding to promoters of mitochondrial-related nuclear gene in NAc. **(A)** Illustration of the ChIP assay using anti-Egr3 to immunoprecipitate chromatin in NAc tissue. **(B) Top panel:** The timeline of cocaine (20mg/kg, i.p.) or saline injections and NAc tissue collection after 24-hour abstinence. **Bottom panel:** Egr3 binding is increased on *Drp1, Nrf2* and *Polγ* gene promoters in NAc in the cocaine group (n=35 mice, 7 samples) compared to saline injected in male (n=30, 6 samples) mice. Student’s t test, *p<0.05; Error bars, SEM.

Next, qRT-PCR was used to quantify mitochondria-related transcripts in bulk NAc tissue to determine if a history of cocaine exposure alters the expression of the following mitochondria-related genes: *Nrf1, Nrf2, PGC1α, Polγ, Tfam, Tfb1*, and *Tomm20*. Mice receiving 7 days of i.p. cocaine (20mg/kg) injections showed an increase in *Nrf1, Nrf2*, and *Tomm20* (see Fig 2a & Additional File S3; for mouse *PGC1*α data, please see Ramesh Chandra, Engeln, Francis, et al., 2017). Rats that self-administered cocaine for 10 days (1mg/kg/infusion) showed similar patterns, with enhancements in *Nrf1, Tfb1*, and *Tomm20* (see Fig 2b & Additional File S3). To compare rodent models with cocaine-dependent individuals, postmortem human tissue was analyzed for the same mitochondria-related nuclear genes. Tissue from cocaine-dependent individuals had enhanced *Nrf2* and *PGC1*α (see Fig 2c & Additional File S3), which is consistent with previous studies demonstrating increased Drp1 in NAc of the same cocaine dependent cohort (1).

**Figure 2.**
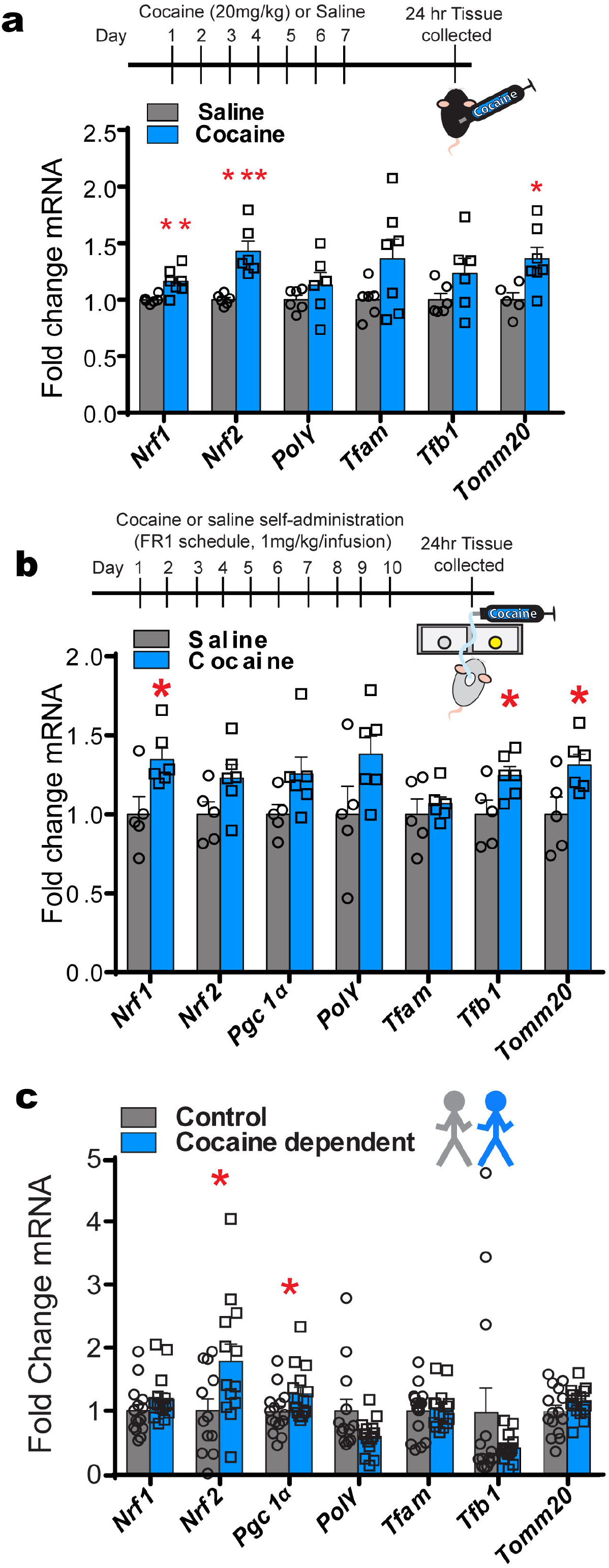
Cocaine induced mitochondrial-related nuclear gene expression in NAc of rodents and human subjects. **(A)** Male mice treated with 7 days of cocaine (20mg/kg, i.p.) display higher mRNA expression of *Nrf1, Nrf2*, and *Tomm20* genes 24h in NAc. Student’s t-test, *p<0.05, ***p<0.001; n=6 saline and n= 7 cocaine group. **(B)** Male rat mRNA expression in NAc tissue after 10 days self-administration **(upper panel)** followed by 24hr abstinence. **Bottom panel**: increased expression of *Nrf1, Tfb1* and *Tomm20* in NAc of rats that self-administer cocaine compared to saline. Student’s t test, *p<0.05; n=5 saline and n= 6 cocaine. **(C)** mRNA expression in human postmortem NAc tissue indicated that *Nrf2* and *PGC1*α gene expression is higher in human cocaine dependents compared to control subjects. Student’s t test, *p<0.05; n=15 control (14 male; 1 female) and n= 15cocaine dependent (14 male;1 female); Error bars, SEM.

While cocaine-induced changes were observed in bulk tissue, previous work has shown differential cocaine-induced regulation of Egr3, the mitochondrial biogenesis genes, Drp1 and PGC1α, and mitochondrial morphology in D1 vs D2-MSNs (1,2,14,39). Egr3, Drp1, and PGC1α are oppositely regulated in D1-MSNs vs D2-MSNs following repeated cocaine exposure (14), and mitochondrial morphology at both baseline and after repeated cocaine is different between D1 and D2-MSNs (1,2,39). Thus, next examined were mitochondrial-related nuclear genes in D1-MSNs vs D2-MSNs using D1-Cre-RT or D2-Cre-RT mice (Fig 3a). D1-Cre-RT mice receiving 7 days of i.p. cocaine injections (20mg/kg) showed an increase in *Tfam* mRNA in NAc D1-MSNs (Fig 3b). By contrast, D2-Cre-RT mice showed a reduction in mRNA levels of *Polγ Tfam, Tfb1*, and *Tomm20* in NAc D2-MSNs after repeated cocaine (Fig 3c). These data are consistent with previous studies demonstrating enhanced Drp1 and PGC1α in NAc D1-MSNs and reduced in D2-MSN after repeated cocaine exposure (2).

**Figure 3.**
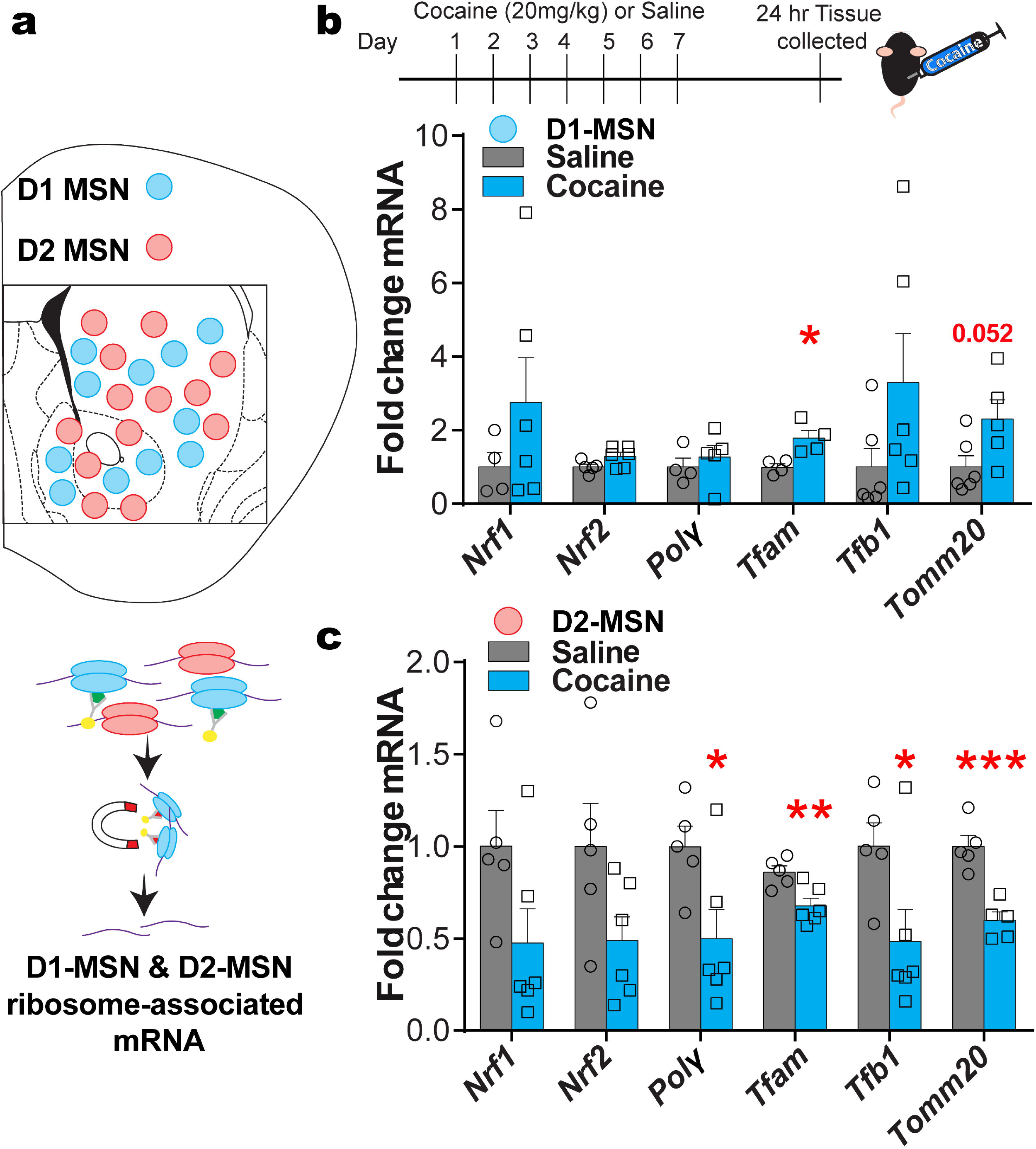
Ribosome associated mitochondrial-related mRNA are altered in NAc MSN subtypes after repeated cocaine. **(A)** Illustration of NAc D1-MSN (blue) and D2-MSN (red) subtypes and the RiboTag procedure to isolate ribosome-associated mRNA from MSN subtypes using D1-Cre-RT and D2-Cre-RT mice. HA tagged (green) ribosomes are immunoprecipitated from NAc homogenates using anti-HA coupled to magnetic beads, followed by isolation of ribosome-associated mRNA from D1-MSNs and D2-MSNs. **(B) Top panel**: The timeline of cocaine (20mg/kg, i.p.) or saline injections and NAc tissue collection after 24-hour abstinence. **Bottom panel**: *Tfam* ribosome-associated mRNA is increased in D1-MSNs Student’s t-test, *p<0.05; p=0.07 *Nrf2;* p=0.052 *Tomm20;* D1-MSN saline group (n=16 females, 8 male; pooled samples 4 and 2, respectively); D1-MSN cocaine group (n=8 female, 16 males; pooled samples 2 and 4, respectively). **(C)** *Polγ, Tfam, Tfb1* and *Tomm20* expression is reduced in D2-MSNs after repeated cocaine (7 days, 20mg/kg). Student’s t-test, *p<0.05; **p<0.01;***p<0.001; D2-MSN saline group (n=8 female, 16 male; pooled samples 2 and 4, respectively) and D2-MSN cocaine group (n=12 female, n=12 male; pooled samples 3 and 3, respectively); Error bars, SEM.

Next we examined whether reducing Egr3 in D1-MSNs alters repeated cocaine-induced expression of mRNAs in this neuron subtype. D1-Cre-RT mice received NAc injections of AAV-DIO-Egr3-miR or AAV-DIO-SS-miR followed by repeated cocaine (20mg/kg for 7 days) or saline followed by 24-hour abstinence (Fig 4a). *PGC1*α and *Drp1* mRNA in particular were analyzed since these were the only two molecules that displayed both an increase in Egr3 binding at promoters and increased mRNA in D1-MSNs after repeated cocaine exposure in this current study or in our previous studies (1,2). D1-Cre-RT mice with Egr3-miR injections into NAc showed 5-fold lower *PGC1*α mRNA and 6-fold lower *Drp1* mRNA (Fig 4b) in D1-MSNs following cocaine injections compared to SS-miR controls that received cocaine. Egr3-miR cocaine mice also showed no difference compared to Egr3-miR saline controls in either *PGC1*α or *Drp1* mRNA. These findings imply Egr3 transcriptionally regulates mitochondrial-related nuclear genes and is consistent with increased Egr3 binding on promoters of Drp1 (Fig 1b) and PGC1α following cocaine exposure (2).

**Figure 4.**
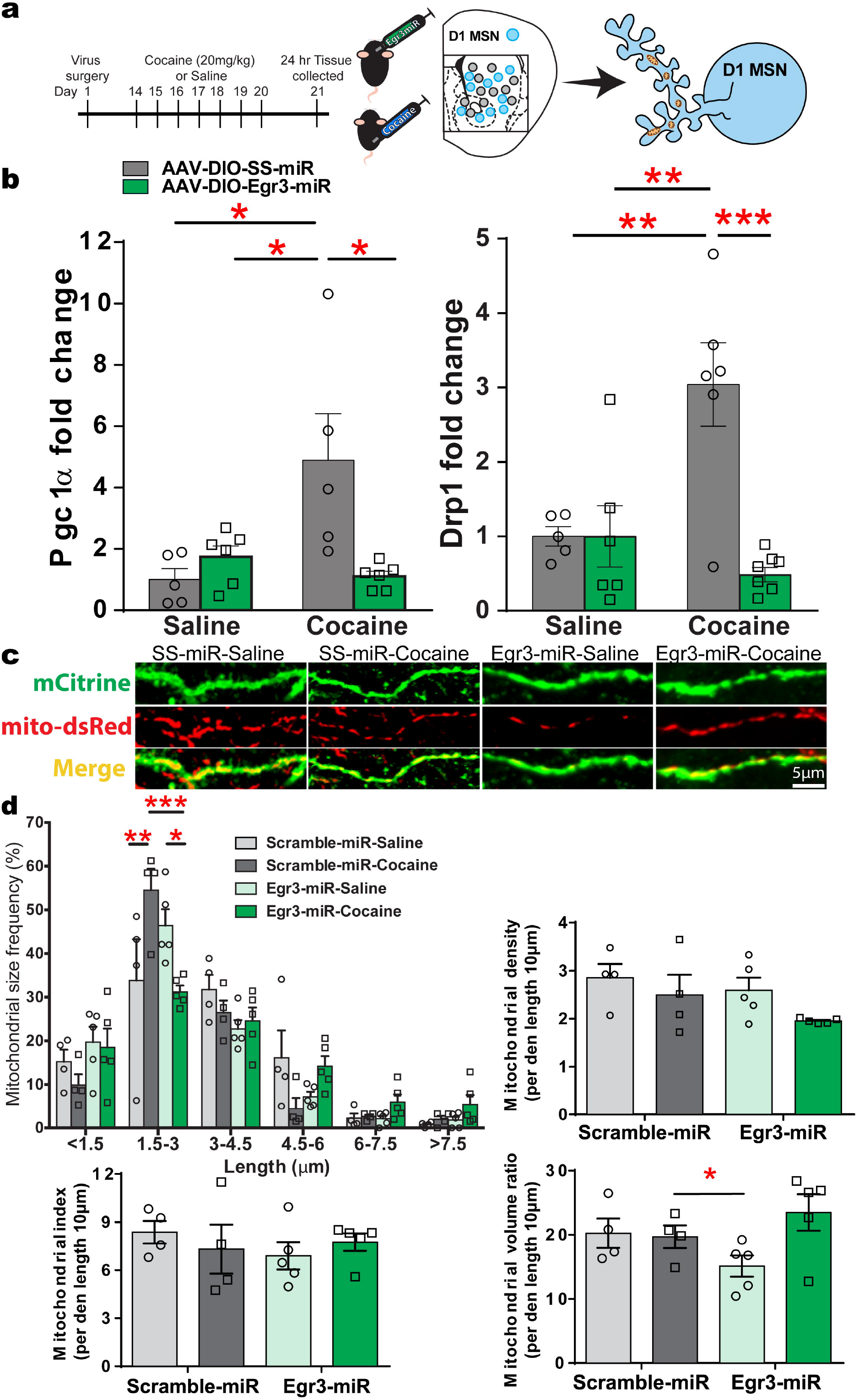
Mitochondria-related nuclear gene expression in NAc D1-MSNs and attenuation of cocaine-mediated induction of small sized mitochondria after Egr3 knockdown. **(A)** D1-Cre-RT mice received an intracranial infusion of a Cre-dependent virus inducing D1-MSN functional knockdown via Egr3-miR (experimental) or SS-miR (control). The timeline of Egr3-miR virus surgery followed by cocaine (20mg/kg, i.p.) or saline injections and NAc tissue collection after 24-hour abstinence. **B)** *Drp1* and *PGC1*α mRNAs are reduced in D1-MSN of the Egr3miR cocaine treated compared to SS-miR cocaine treated male mice. Two-way ANOVA: Drp1; Interaction F_(1,20)_= 12.67; Post hoc test p = 0.002, **p<0.01;***, p<0.001; SS-miR Saline (n=3 male, n=2 female), SS-miR Cocaine (n=3 female, n=3 male), Egr3-miR Saline (n=4 male, n=2 female), Egr3-miR cocaine (n=3 female, n=4 male). Two-way ANOVA: PGC1α; Interaction F_(1,18)_= 9.398; Post hoc test p= 0.0067, *p<0.05; n=5 SS-miR Saline, n=5 SS-miR Cocaine, n=6 Egr3-miR Saline, n=6 Egr3-miR cocaine all males. **(C)** Representative images for D1-MSN dendrites (green) with labeled mitochondria (red) after repeated cocaine exposure from SS-miR and Egr3-miR groups. **(D) Top left:** 3D reconstruction of dendrites and mitochondria highlights an increase in the frequency of smaller length mitochondria in D1-MSN dendrites in the cocaine SS-miR group, which is attenuated in the Egr3-miR cocaine group. The Egr3-miR cocaine group displayed a decrease in the frequency of smaller size mitochondria in the 1.5-3μm size bracket compared to the SS-miR cocaine group. Two-way ANOVA: Interaction: F_(15,70)_= 3.035, p < 0.0001, Bonferroni post hoc: *p<0.05,**p<0.01;***p<0.001; SS-miR saline (n=4, all male); SS-miR cocaine (n=4, all male); Egr3-miR saline (n=2 female, n=3 male); Egr3-miR cocaine (n=2 female, n=3 male). **Top Right:** Mitochondrial density (per dendrite 10μm length) is unaltered in D1-MSNs of mice exposed to cocaine and Egr3-miR. Two-way ANOVA:F _(1,14)_= 0.2823; p= 0.6035;. **Bottom Left:** Mitochondrial index (mitochondrial length per 10μm of dendrite length) in Egr3 knockdown compared to control group demonstrating no change among the four groups. Two-way ANOVA F _(1,14)_ = 1.041;p= 0.8304;. **Bottom Right:** Mitochondrial volume ratio (relative to dendrite volume) is enhanced in the SS-miR group compared to the Egr3-miRgroup. Two-way ANOVA F_(1,14)_=3.919;p= 0.0678;;Error bars, SEM.

Drp1 mediates enhanced mitochondrial fission in D1-MSN dendrites following repeated cocaine exposure, including self-administration and blockade of Drp1 prevents fission in these neurons. (1). Next examined was the impact of Egr3 reduction on cocaine-induced mitochondrial fission. D1-Cre mice received NAc AAV-DIO-Egr3-miR or AAV-DIO-SS-miR along with expressing AAV-DIO-mito-dsRed (1,39) to label mitochondria in D1-MSNs, and to allow quantification of mitochondrial volume, density, and size (Fig 4d). Mice received 7 days of i.p. cocaine injections (20mg/kg) or saline, and 24-hours after the last injection brain tissue was collected. An increase in the frequency of smaller-length mitochondria in dendrites was observed in cocaine treated mice, relative to saline treated mice, consistent with previous findings (Fig 4d; Ramesh Chandra, Engeln, Schiefer, et al., 2017). However, the frequency of small-sized mitochondria was decreased by approximately 30% in Egr3-miR cocaine exposed mice relative to Egr3-miR saline controls. SS-miR controls treated with cocaine showed the highest frequency in the 1.5-3.0μm length, whereas Egr3-miR mice treated with cocaine showed about half as many of these small-sized mitochondria (Two-way ANOVA: Interaction: F_(15,70)_= 3.035, p < 0.0001, Bonferroni post hoc: *p<0.05,**p<0.01;***p<0.001;). By contrast, Egr3-miR mice treated with cocaine showed the most large-sized mitochondria in the 4.5-6μm length. We observed no changes in mitochondrial density or index by both drug and virus conditions. However, Egr3-miR mice demonstrated a lower volume than SS-miR mice. (Fig 4d; Two-way ANOVA F_(1,14)_=3.919;p= 0.0678; Error bars, SEM.). Taken together, cocaine-induced gene regulation and mitochondrial structural characteristics are substantially altered in NAc D1-MSNs. These findings align with previous work showing that manipulation of Drp1 dictates mitochondrial morphology characteristics following cocaine self-administration (1).

## Discussion

Cocaine alters NAc mitochondrial morphology, inducing small-sized mitochondria in D1-MSNs and shifts toward larger mitochondria in D2-MSNs (1). This study sought to identify relationships between the cocaine-responsive transcription factor, Egr3, and genes regulating mitochondrial machinery to confirm causal mechanistic connections between Egr3 regulation and mitochondrial dynamics and morphology characteristics. This study identified distinct gene expression patterns of mitochondrial-related genes after cocaine exposure in NAc and in subpopulations of D1 and D2-MSNs. Egr3 was shown as an upstream transcriptional regulator of a number of mitochondrial nuclear genes following cocaine exposure. Further, this study established causal impacts of Egr3 regulation on mitochondria-related genes and the functional impacts of gene regulation on mitochondrial shape following cocaine exposure in NAc D1-MSNs. Together, the presented findings are some of the first examinations of NAc and neuron subpopulation responses to cocaine for a number of mitochondrial-associated factors regulating mitochondrial machinery and subsequent morphological profiles.

First, repeated cocaine exposure upregulated Egr3 binding to mitochondria-related nuclear gene promoters in NAc, aligning with previous findings showing enhanced Egr3 binding to PGC1α following cocaine exposure (2). Subsequent tests examined mRNA of Egr3-target mitochondrial nuclear genes in NAc. These assays revealed that mice receiving cocaine injections and rats self-administering cocaine showed similar patterns of gene expression, with enhanced *Nrf1, Nrf2*, and *Tomm20*. However, only *PGC1*α and *Nrf2* were upregulated in postmortem tissue of cocaine dependents. Additionally, other studies have shown that cocaine upregulates the mitochondrial fission genes Drp1 (1,2) in bulk NAc tissue; including cocaine enhancement of *Drp1* mRNA in both rodents and humans (1). The lack of *PGC1*α enhancement in NAc bulk tissue of rats that self-administer cocaine is inconsistent to the increases observed in postmortem NAc of cocaine dependents and NAc D1-MSNs in mice exposed to cocaine. This could be due to masked cell subtype changes since the analysis was performed in rat bulk NAc tissue.

What might account for the disparity in gene expression between rodents and humans? Factors, such as route of cocaine administration, methods of extraction, and potential co-morbid mental disorders in human subjects, such as major depression or bipolar disorder. These differences may reflect neurobiological differences in genotype and phenotype regulation patterns in the NAc and in MSN subtypes linked to broad forms of neuropathology (9–11,45–47).

Analysis of NAc-MSN subtype mRNA demonstrated upregulation of mitochondria-related mRNAs in D1-MSNs, whereas D2-MSNs displayed a general decrease in mitochondria-related mRNAs. In comparison to bulk NAc tissue quantification, these findings again demonstrate the importance of localizing and characterizing gene regulation to specific neuron subtypes within regions of interest. Further, these findings are consistent with a previous study demonstrating increases in Drp1 expression and increased frequency of small mitochondria in D1-MSNs, while lower levels of Drp1 and increased larger mitochondria frequency are observed in D2-MSNs following cocaine exposure or self-administration (1). Moreover, PGC1α shows enhancement in D1-MSNs and a decrease in D2-MSNs following 7 days of cocaine administration (2). Thus, these findings further elucidate the differences of mitochondrial-related molecules in MSN subtypes following repeated cocaine exposure. The different patterns of mitochondrial nuclear gene regulation align with previous work showing D1-MSN specific enhancement of Egr3, PGC1α, and Drp1 following cocaine exposure (1,2). Collectively, differences in regulation between these neuron populations highlight an important principle: D1-MSNs vs D2-MSNs differentially regulate mitochondrial nuclear genes in response to cocaine. Subsequent disruption of Egr3 by NAc infusion of Egr3-miR into D1-MSNs prevented cocaine-mediated enhancement of *Drp1* and *PGC1*α mRNAs. These findings corroborate a model of Egr3 transcriptional regulation of mitochondrial nuclear genes, and previous findings showing both cocaine-enhanced Egr3-PGC1α binding and cocaine-induced upregulation of PGC1α and Drp1 in D1 MSNs. (1,2,14).

The importance of examining cell type-specific changes is further highlighted by cocaine induction of Egr3 promoter binding to Drp1 and the prevention of Drp1 upregulation in D1-MSNs by Egr3 blockade; indicating Egr3 transcriptional regulation of Drp1 is likely necessary for D1-MSN Drp1-mediated mitochondrial fission after cocaine exposure. D1-MSNs are known to have enhanced spine density after cocaine exposure (20,25) and Drp1 is necessary for the formation of dendritic spines (23,24). Here, Egr3 blockade in D1-MSNs also prevented cocaine-induced fission, indicated by reduced frequency of small-sized mitochondria. Thus, converging lines of evidence support the hypothesis that cocaine-induced neuroplasticity in NAc, particularly in NAc D1-MSNs, is directly tuned by Egr3 regulation of mitochondrial dynamics. Given a large literature demonstrating enhanced excitatory plasticity, signaling processes, and transcriptional activity in D1-MSNs with both contingent and non-contingent chronic or acute cocaine it is plausible that these neurons require higher energy demands (13,17–20,48,49) or mitochondria could be acting as buffers of calcium homeostasis required for neuroplastic events (23). Thus, cocaine-induced enhancement of Egr3 and subsequent mitochondrial fission may work to meet increased energy demands. Enhancement of small-sized mitochondria indicates enhanced fission; here impaired by Egr3-miR disruption of Egr3 function. The alterations of mitochondrial morphology observed here may be responsible for facilitating long-lasting neuroplastic changes underpinning persistent craving of drugs of abuse.

Egr3 is necessary for cocaine-induced enhancement of Drp1, cocaine self-administration produces morphological shifts to small-sized mitochondria in D1-MSNs, and Drp1 and fission are required for neuroplasticity in neurons including after cocaine exposure (1,23). It is plausible that the alterations of the other mitochondria nuclear genes affected by cocaine and Egr3 in this study are involved in these dynamic changes. Egr3-PGC1α enhancement also indicates recruitment of upstream fission/biogenesis processes and energy production machinery (2). PGC1α knockdown results in decreased neurofilament protein (22) and PGC1α is involved in the formation and maintenance of dendritic spines (50). Thus, PGC1α may be critical for the dendritic spine and corresponding plasticity adaptations occurring with cocaine exposure(43,51). Further, PGC1α null mice show impairment in sodium ion pumps (22) and neurons require large quantities of ATP to maintain ion gradients and transport mechanisms. While PGC1α is involved in energy production it is also involved in the regulation of reactive oxygen species (ROS) production, a byproduct of cellular respiration (52,53). PGC1α regulates factors in the detoxification of ROS, such as superoxide dismutase, indicating concordant cell growth and protective effects by both enhancing cellular respiration and scavenging of toxic byproduct (54). This suggests a neuroprotective effect from PGC1α regulation.

Egr3 also regulates CREB, a cocaine-responsive gene, which in turn regulates several mitochondrial components (R. Chandra et al., 2015). For instance, CREB regulates the expression of PGC1α (55,56). CREB directly regulates cytochrome c (a major factor in ETC function), *PGC1*, and *Nrf1* (55,57–62). Further, tandem CREB-Nrf1 activity is involved in the growth-regulated cytochrome c (57,60). CREB is also mediated by TOM complexes into mitochondria, allowing access to and binding on mtDNA (58,63,64). This activity of nuclear CREB and mitochondrial CREB is coordinated via shared upstream signals. Taken together, Egr3 or other mitochondrial gene regulation induced by cocaine or viral manipulations may impact mitochondrial function indirectly through CREB activity on mitochondria-related nuclear genes to facilitate neuronal adaptation.

The result of low sample size may present false negatives in this report and hide cocaine-induced effects, such as in the case of ChIP assaying of Tfam and Tfb1 in Fig 1. However, it is noted for ChIP and RiboTag assays that samples displayed are representative of pooled tissues from 5 or 4 mice, respectively, and, thus, likely representative of authentic effects.

While this study establishes a causal connection between D1-MSN Egr3 expression and mitochondrial fission, a limitation still to be addressed is how D1-MSN Egr3-mediated mitochondrial fission affects cocaine-induced behavioral adaptations. As discussed above, studies of the fission-regulating molecules Drp1 and PGC1α show that enhancement of D1-MSN mitochondrial fission increases some cocaine-induced behaviors, such as seeking for cocaine, cocaine CPP, and cocaine-induced locomotion (1,2). Thus, because cocaine exposure enhances Egr3 in D1-MSNs (14), enhances Egr3 binding to PGC1α (2), and is shown here to induce enhanced Egr3-Drp1 binding, it is predicted that subsequent enhancements of D1-MSN mitochondrial fission would induce increased cocaine-induced behaviors, such as locomotion, CPP or seeking for cocaine. It would be of interest examine whether modulation of fission or fusion-related molecules is capable of combating the effects of Egr3 manipulations. For instance, it is possible that Egr3-miR in D1-MSNs, which decreases mitochondrial fission, would be counteracted by Drp1 overexpression, shown to enhance cocaine seeking, and result in reversal of cocaine-induced behavioral impairment. Similarly, enhanced PGC1α may counter Egr3 knockdown to restore cocaine CPP or locomotion. Alternatively, D1-MSN Drp1 or PGC1α blockade could impair enhancement of cocaine-induced behaviors by Egr3.

A related interesting question is whether enhanced mitochondrial fusion would run counter to D1-MSN Egr3 responses to cocaine. If cocaine induces enhanced D1-MSN Egr3 expression and subsequent mitochondrial fission, it would be expected that overexpression of fusion molecules would produce behavioral effects counter to Egr3.. However, previous studies have examined the expression of the mitochondrial fusion genes, Mitofusin 1 and 2 (Mfn1 and Mfn2) and optic atrophy 1 (Opa1), but found relatively little effect of cocaine on these genes in general NAc cell populations (1). Further, D1 vs D2-MSN characterization of these genes after repeated cocaine injections showed no differences between neuron subpopulations. However, in NAc Mfn2 expression is elevated after cocaine self-administration in rats, but it remains to be seen whether this is driven by specific neuron subpopulations. An outstanding possibility is that cocaine self-administration could induce D1-MSN changes in fusion genes, such as Mfn2, whereas i.p. cocaine does not. Thus, it is conceivable that artificial overexpression of fusion genes could counteract D1-MSN Egr3-mediated fission, no firm opinions can be made given thus far cocaine intake has little impact or can even enhance fusion-related molecules. Ultimately, the answers to how Egr3 regulation of mitochondrial dynamics relate to psychological impact remain to be determined by future studies.

## Conclusion

Together, these findings establish that Egr3 regulates enhancement of mitochondrial dynamics, that have been shown to regulate plasticity adaptations in MSNs underlying behaviors occurring in rodent models of substance use disorders. Further, these studies are some of the first illustrations of cell type-specific gene regulation in brain and specifically the NAc. These studies support a model whereby cocaine-associated transcription factors are controlling mechanisms of mitochondrial dynamics and underpin long-lasting neurobiological adaptations to drugs of abuse.

## List of abbreviations

AAV: adeno-associated viruses
ChIP: chromatin immunoprecipitation
CPP: conditioned place preference
D1-MSN: dopamine receptor-1 containing medium spiny neurons
D2-MSN: dopamine receptor-2 containing medium spiny neurons
DSM-IV: Diagnostic and Statistical Manual of Mental Disorders-IV
DIO: double inverted open
Drp1: dynamin related protein 1
Egr3: early growth response factor 3
Egr3-miR: Egr3 microRNA
IACUC: Institutional Animal Care and Use Committee
Mfn1: Mitofusin 1
Mfn2: Mitofusin 2
MSNs: medium spiny neurons
NAc: Nucleus accumbens
Opa1: Optic atrophy 1 (alternative: OPA1 mitochondrial dynamin like GTPase)
PBS: phosphate buffered saline
PGC1α: peroxisome proliferator-activated receptor gamma coactivator
Polγ: mitochondrial DNA polymerase subunit gamma
Nrf1: transcription factors nuclear respiratory factor 1
Nrf2: transcription factors nuclear respiratory factor 2
qRT-PCR: quantitative real time polymerase chain reaction
RT: RiboTag
SS-miR: scramble sequence microRNA
Tfam: mitochondria-specific transcription factor a
Tfb1: mitochondria-specific transcription factor b
Tomm20: translocase of the outer mitochondrial membrane 20

## Author Contribution

All authors contributed to the manuscript. All authors have given final approval for this version to be published and agree to be accountable for aspects of the work:

S.C., R.C., M.H., I.P., T.W., H.K., L.J., and A.M.G. performed experiments. G.T. and S.J.R. provided and processed postmortem human cDNA. S.C., R.C., A.M.G., D.M.D., and M.K.L designed experiments. S.C., R.C., and M.K.L. wrote and edited the manuscript.

## Acknowledgements

The authors would like to thank N.G. Larsson (Max Planck Institute for Biology of Ageing, D-50931 Cologne, Germany) for supplying the mito-dsRed vector used in this study.

---

**Additional File 1. Human demographics for examined cocaine dependent and non-dependent tissue**

**Additional File 2. Primers utilized in study**

**Additional File 3. Gene regulation by species, brain region, and cell population**

**Additional File 4. Statistical table**

## References

1. Chandra R, Engeln M, Schiefer C, Patton MH, Martin JA, Werner CT, et al. Drp1 Mitochondrial Fission in D1 Neurons Mediates Behavioral and Cellular Plasticity during Early Cocaine Abstinence. Neuron. 2017;96(6):1327–1341.e6.

2. Chandra R, Engeln M, Francis TC, Konkalmatt P, Patel D, Lobo MK. A Role for Peroxisome Proliferator-Activated Receptor Gamma Coactivator-1alpha in Nucleus Accumbens Neuron Subtypes in Cocaine Action. Biol Psychiatry. 2017 Apr;81(7):564–72.

3. Cunha-Oliveira T, Silva L, Silva AM, Moreno AJ, Oliveira CR, Santos MS. Mitochondrial complex I dysfunction induced by cocaine and cocaine plus morphine in brain and liver mitochondria. Toxicol Lett. 2013 Jun;219(3):298–306.

4. Volkow ND, Koob GF, McLellan AT. Neurobiologic Advances from the Brain Disease Model of Addiction. N Engl J Med [Internet]. 2016;374(4):363–71. Available from: https://www.ncbi.nlm.nih.gov/pubmed/26816013

5. Baumgartner HM, Cole SL, Olney JJ, Berridge KC. Desire or Dread from Nucleus Accumbens Inhibitions: Reversed by Same-Site Optogenetic Excitations. J Neurosci. 2020;

6. Gerfen CR, Engber TM, Mahan LC, Susel Z, Chase TN, Monsma Jr. FJ, et al. D1 and D2 dopamine receptor-regulated gene expression of striatonigral and striatopallidal neurons. Science (80-). 1990/12/07. 1990;250(4986):1429–32.

7. Smith RJ, Lobo MK, Spencer S, Kalivas PW. Cocaine-induced adaptations in D1 and D2 accumbens projection neurons (a dichotomy not necessarily synonymous with direct and indirect pathways). Curr Opin Neurobiol [Internet]. 2013/02/18. 2013 Aug;23(4):546–52. Available from: https://pubmed.ncbi.nlm.nih.gov/23428656

8. Fox ME, Chandra R, Menken MS, Larkin EJ, Nam H, Engeln M, et al. Dendritic remodeling of D1 neurons by RhoA/Rho-kinase mediates depression-like behavior. Mol Psychiatry. 2018 Aug;

9. Francis TC, Chandra R, Friend DM, Finkel E, Dayrit G, Miranda J, et al. Nucleus accumbens medium spiny neuron subtypes mediate depression-related outcomes to social defeat stress. Biol Psychiatry. 2015;

10. Massaly N, Copits BA, Wilson-Poe AR, Hipólito L, Markovic T, Yoon HJ, et al. Pain-Induced Negative Affect Is Mediated via Recruitment of The Nucleus Accumbens Kappa Opioid System. Neuron. 2019;

11. Ren W, Centeno MV, Berger S, Wu Y, Na X, Liu X, et al. The indirect pathway of the nucleus accumbens shell amplifies neuropathic pain. Nat Neurosci [Internet]. 2016;19(2):220–2. Available from: https://doi.org/10.1038/nn.4199

12. Lobo MK, Covington HE 3rd, Chaudhury D, Friedman AK, Sun H, Damez-Werno D, et al. Cell type-specific loss of BDNF signaling mimics optogenetic control of cocaine reward. Science (80-). 2010/10/16. 2010;330(6002):385–90.

13. Lobo MK, Zaman S, Damez-Werno DM, Koo JW, Bagot RC, DiNieri JA, et al. DeltaFosB induction in striatal medium spiny neuron subtypes in response to chronic pharmacological, emotional, and optogenetic stimuli. J Neurosci. 2013 Nov;33(47):18381–95.

14. Chandra R, Francis TC, Konkalmatt P, Amgalan A, Gancarz AM, Dietz DM, et al. Opposing Role for Egr3 in Nucleus Accumbens Cell Subtypes in Cocaine Action. J Neurosci [Internet]. 2015;35(20):7927–37. Available from: http://www.jneurosci.org/cgi/doi/10.1523/JNEUROSCI.0548-15.2015

15. Bertran-Gonzalez J, Bosch C, Maroteaux M, Matamales M, Herve D, Valjent E, et al. Opposing patterns of signaling activation in dopamine D1 and D2 receptor-expressing striatal neurons in response to cocaine and haloperidol. J Neurosci. 2008/05/30. 2008;28(22):5671–85.

16. Bock R, Shin JH, Kaplan AR, Dobi A, Markey E, Kramer PF, et al. Strengthening the accumbal indirect pathway promotes resilience to compulsive cocaine use. Nat Neurosci. 2013 May;16(5):632–8.

17. Lobo MK, Nestler EJ. The striatal balancing act in drug addiction: distinct roles of direct and indirect pathway medium spiny neurons. Front Neuroanat. 2011;5:41.

18. Grueter BA, Robison AJ, Neve RL, Nestler EJ, Malenka RC. FosB differentially modulates nucleus accumbens direct and indirect pathway function. Proc Natl Acad Sci U S A. 2013 Jan;110(5):1923–8.

19. MacAskill AF, Cassel JM, Carter AG. Cocaine exposure reorganizes cell type- and input-specific connectivity in the nucleus accumbens. Nat Neurosci. 2014 Sep;17(9):1198–207.

20. Kim J, Park BH, Lee JH, Park SK, Kim JH. Cell type-specific alterations in the nucleus accumbens by repeated exposures to cocaine. Biol Psychiatry [Internet]. 2011;69(11):1026–34. Available from: http://dx.doi.org/10.1016/j.biopsych.2011.01.013

21. Finck BN, Kelly DP. PGC-1 coactivators: inducible regulators of energy metabolism in health and disease. J Clin Invest. 2006 Mar;116(3):615–22.

22. Lin J, Handschin C, Spiegelman BM. Metabolic control through the PGC-1 family of transcription coactivators. Cell Metab. 2005 Jun;1(6):361–70.

23. Divakaruni SS, Van Dyke AM, Chandra R, LeGates TA, Contreras M, Dharmasri PA, et al. Long-Term Potentiation Requires a Rapid Burst of Dendritic Mitochondrial Fission during Induction. Neuron [Internet]. 2018;1–16. Available from: https://linkinghub.elsevier.com/retrieve/pii/S0896627318308274

24. Li Z, Okamoto KI, Hayashi Y, Sheng M. The importance of dendritic mitochondria in the morphogenesis and plasticity of spines and synapses. Cell. 2004;119(6):873–87.

25. Bobadilla A-C, Heinsbroek JA, Gipson CD, Griffin WC, Fowler CD, Kenny PJ, et al. Corticostriatal plasticity, neuronal ensembles, and regulation of drug-seeking behavior. Prog Brain Res. 2017;235:93–112.

26. Cahill ME, Walker DM, Gancarz AM, Wang ZJ, Lardner CK, Bagot RC, et al. The dendritic spine morphogenic effects of repeated cocaine use occur through the regulation of serum response factor signaling. Mol Psychiatry [Internet]. 2017 May 30;23:1474. Available from: https://doi.org/10.1038/mp.2017.116

27. Wolf ME. Synaptic mechanisms underlying persistent cocaine craving. Nat Rev Neurosci. 2016 Jun;17(6):351–65.

28. Ferrario CR, Gorny G, Crombag HS, Li Y, Kolb B, Robinson TE. Neural and behavioral plasticity associated with the transition from controlled to escalated cocaine use. Biol Psychiatry. 2005 Nov;58(9):751–9.

29. Gipson CD, Kupchik YM, Kalivas PW. Rapid, transient synaptic plasticity in addiction. Neuropharmacology [Internet]. 2014;76(PART B):276–86. Available from: http://dx.doi.org/10.1016/j.neuropharm.2013.04.032

30. Norrholm SD, Bibb JA, Nestler EJ, Ouimet CC, Taylor JR, Greengard P. Cocaine-induced proliferation of dendritic spines in nucleus accumbens is dependent on the activity of cyclin-dependent kinase-5. Neuroscience. 2003/01/22. 2003;116(1):19–22.

31. Robinson TE, Kolb B. Structural plasticity associated with exposure to drugs of abuse. Neuropharmacology. 2004;47 Suppl 1:33–46.

32. Wang X, Cahill ME, Werner CT, Christoffel DJ, Golden SA, Xie Z, et al. Kalirin-7 mediates cocaine-induced AMPA receptor and spine plasticity, enabling incentive sensitization. J Neurosci. 2013 Jul;33(27):11012–22.

33. Gancarz AM, Wang ZJ, Schroeder GL, Damez-Werno D, Braunscheidel KM, Mueller LE, et al. Activin receptor signaling regulates cocaine-primed behavioral and morphological plasticity. Nat Neurosci. 2015;18(7):959–61.

34. Engeln M, Mitra S, Chandra R, Gyawali U, Fox ME, Dietz DM, et al. Sex-Specific Role for Egr3 in Nucleus Accumbens D2-Medium Spiny Neurons Following Long-Term Abstinence From Cocaine Self-administration. Biol Psychiatry. 2020 Jun;87(11):992–1000.

35. Gerfen CR, Paletzki R, Heintz N. GENSAT BAC cre-recombinase driver lines to study the functional organization of cerebral cortical and basal ganglia circuits. Neuron. 2013;

36. Chandra R, Lobo MK. Beyond Neuronal Activity Markers: Select Immediate Early Genes in Striatal Neuron Subtypes Functionally Mediate Psychostimulant Addiction. Front Behav Neurosci. 2017;11:112.

37. Golden SA, Christoffel DJ, Heshmati M, Hodes GE, Magida J, Davis K, et al. Epigenetic regulation of RAC1 induces synaptic remodeling in stress disorders and depression. Nat Med. 2013 Mar;19(3):337–44.

38. Robison AJ, Vialou V, Mazei-Robison M, Feng J, Kourrich S, Collins M, et al. Behavioral and structural responses to chronic cocaine require a feedforward loop involving ΔFosB and calcium/calmodulin-dependent protein kinase II in the nucleus accumbens shell. J Neurosci. 2013 Mar;33(10):4295–307.

39. Chandra R, Calarco CA, Lobo MK. Differential mitochondrial morphology in ventral striatal projection neuron subtypes. J Neurosci Res. 2019 Dec;97(12):1579–89.

40. Chandra R, Lenz JD, Gancarz AM, Chaudhury D, Schroeder GL, Han M-H, et al. Optogenetic inhibition of D1R containing nucleus accumbens neurons alters cocaine-mediated regulation of Tiam1. Front Mol Neurosci. 2013;6:13.

41. Maze I, Covington HE 3rd, Dietz DM, LaPlant Q, Renthal W, Russo SJ, et al. Essential role of the histone methyltransferase G9a in cocaine-induced plasticity. Science. 2010 Jan;327(5962):213–6.

42. Renthal W, Kumar A, Xiao G, Wilkinson M, Covington HE, Maze I, et al. Genome-wide Analysis of Chromatin Regulation by Cocaine Reveals a Role for Sirtuins. Neuron. 2009;

43. Russo SJ, Dietz DM, Dumitriu D, Morrison JH, Malenka RC, Nestler EJ. The addicted synapse: mechanisms of synaptic and structural plasticity in nucleus accumbens. Trends Neurosci. 2010/03/09. 2010;33(6):267–76.

44. Feng J, Wilkinson M, Liu X, Purushothaman I, Ferguson D, Vialou V, et al. Chronic cocaine-regulated epigenomic changes in mouse nucleus accumbens. Genome Biol [Internet]. 2014;15(4):R65. Available from: https://doi.org/10.1186/gb-2014-15-4-r65

45. Chen YH, Huang EYK, Kuo TT, Miller J, Chiang YH, Hoffer BJ. Impact of traumatic brain injury on dopaminergic transmission. Cell Transplantation. 2017.

46. Francis TC, Chandra R, Gaynor A, Konkalmatt P, Metzbower SR, Evans B, et al. Molecular basis of dendritic atrophy and activity in stress susceptibility. Mol Psychiatry [Internet]. 2017 Sep 12;22:1512. Available from: https://doi.org/10.1038/mp.2017.178

47. Galvan L, André VM, Wang EA, Cepeda C, Levine MS. Functional Differences Between Direct and Indirect Striatal Output Pathways in Huntington’s Disease. J Huntingtons Dis. 2012;1:17–25.

48. Pascoli V, Terrier J, Espallergues J, Valjent E, O’Connor EC, Luscher C. Contrasting forms of cocaine-evoked plasticity control components of relapse. Nature. 2014 May;509(7501):459–64.

49. Savell KE, Tuscher JJ, Zipperly ME, Duke CG, Phillips RA, Bauman AJ, et al. A dopamine-induced gene expression signature regulates neuronal function and cocaine response. Sci Adv [Internet]. 2020 Jun 1;6(26):eaba4221. Available from: http://advances.sciencemag.org/content/6/26/eaba4221.abstract

50. Cheng A, Wan R, Yang J-L, Kamimura N, Son TG, Ouyang X, et al. Involvement of PGC-1α in the formation and maintenance of neuronal dendritic spines. Nat Commun. 2012;3:1250.

51. Dong Y, Nestler EJ. The neural rejuvenation hypothesis of cocaine addiction. Trends Pharmacol Sci. 2014 Aug;35(8):374–83.

52. Lenaz G, Bovina C, D’Aurelio M, Fato R, Formiggini G, Genova ML, et al. Role of mitochondria in oxidative stress and aging. In: Annals of the New York Academy of Sciences. 2002.

53. Lowell BB, Shulman GI. Mitochondrial dysfunction and type 2 diabetes. Science. 2005.

54. St-Pierre J, Lin J, Krauss S, Tarr PT, Yang R, Newgard CB, et al. Bioenergetic analysis of peroxisome proliferator-activated receptor γ coactivators 1α and 1β (PGC-kt and PGC-1α) in muscle cells. J Biol Chem. 2003;

55. Herzig S, Long F, Jhala US, Hedrick S, Quinn R, Bauer A, et al. CREB regulates hepatic gluconeogenesis through the coactivator PGC-1. Nature. 2001;

56. Ojuka EO. Role of calcium and AMP kinase in the regulation of mitochondrial biogenesis and GLUT4 levels in muscle. Proc Nutr Soc. 2004;

57. Herzig RP, Scacco S, Scarpulla RC. Sequential serum-dependent activation of CREB and NRF-1 leads to enhanced mitochondrial respiration through the induction of cytochrome c. J Biol Chem. 2000;

58. Lee J, Kim CH, Simon DK, Aminova LR, Andreyev AY, Kushnareva YE, et al. Mitochondrial cyclic AMP response element-binding protein (CREB) mediates mitochondrial gene expression and neuronal survival. J Biol Chem. 2005;

59. Franko A, Mayer S, Thiel G, Mercy L, Arnould T, Hornig-Do H-T, et al. CREB-1α Is Recruited to and Mediates Upregulation of the Cytochrome c Promoter during Enhanced Mitochondrial Biogenesis Accompanying Skeletal Muscle Differentiation. Mol Cell Biol. 2008;

60. Gopalakrishnan L, Scarpulla RC. Differential regulation of respiratory chain subunits by a CREB-dependent signal transduction pathway. Role of cyclic AMP in cytochrome c and COXIV gene expression. J Biol Chem. 1994;

61. Suliman HB, Sweeney TE, Withers CM, Piantadosi CA. Co-regulation of nuclear respiratory factor-1 by NFκB and CREB links LPS-induced inflammation to mitochondrial biogenesis. J Cell Sci. 2010;

62. Vercauteren K, Pasko RA, Gleyzer N, Marino VM, Scarpulla RC. PGC-1-Related Coactivator: Immediate Early Expression and Characterization of a CREB/NRF-1 Binding Domain Associated with Cytochrome *c* Promoter Occupancy and Respiratory Growth. Mol Cell Biol [Internet]. 2006 Oct 15;26(20):7409 LP–7419. Available from: http://mcb.asm.org/content/26/20/7409.abstract

63. Cammarota M, Paratcha G, Bevilaqua LRM, De Stein ML, Lopez M, Pellegrino De Iraldi A, et al. Cyclic AMP-responsive element binding protein in brain mitochondria. J Neurochem. 1999;

64. De Rasmo D, Signorile A, Roca E, Papa S. CAMP response element-binding protein (CREB) is imported into mitochondria and promotes protein synthesis. FEBS J. 2009;

