## Additional File 1 for "Cocaine-induced neuron subtype mitochondrial dynamics through Egr3 transcriptional regulation"

| SN # | Cocaine Dependence | Brain ID | Cause of Death | Age | Gender | Weight (g) | PH | Refrigeration Delay (hrs) | Comorbid Disorder |
|---|---|---|---|---|---|---|---|---|---|
| 1 | Cocaine dependent | 7 | Accident | 51 | Male | 1428 | 6.73 | 1.5 | Major Depression |
| 2 | Cocaine dependent | 22 | Natural | 39 | Male | 1140 | 6.86 | 27.5 | Depressive NOS |
| 3 | Cocaine dependent | 26 | Suicide | 36 | Male | 1548 | 6.54 | 12 | BiPolar Disorder |
| 4 | Cocaine dependent | 33 | Suicide | 45 | Male | 1430 | 6.57 | 2.75 | Major Depression |
| 5 | Cocaine dependent | 34 | Suicide | 35 | Male | 1425 | 6.81 | 3.5 | |
| 6 | Cocaine dependent | 112 | Suicide | 24 | Male | 1460 | 6.89 | 2.5 | |
| 7 | Cocaine dependent | 120 | Suicide | 48 | Male | 1460 | 6.56 | 1.75 | Major Depression |
| 8 | Cocaine dependent | 121 | Suicide | 33 | Male | 1580 | 6.75 | 9 | |
| 9 | Cocaine dependent | 140 | Suicide | 43 | Male | 1445 | 6.78 | 5.25 | Psychotic Disorder NOS |
| 10 | Cocaine dependent | 147 | Accident | 24 | Male | 1480 | 6.33 | 7 | |
| 11 | Cocaine dependent | 156 | Suicide | 39 | Male | 1552 | 6.7 | 4.25 | Major Depression |
| 12 | Cocaine dependent | 193 | Suicide | 38 | Male | 1511.4 | 6.5 | 16 | Depressive NOS |
| 13 | Cocaine dependent | 194 | Suicide | 53 | Male | 1366.1 | 6.5 | 28 | |
| 14 | Cocaine dependent | 176 | Suicide | 50 | Female | 1255.5 | 6.3 | 7 | |
| 15 | Not Cocaine Dependent | 128 | Suicide | 46 | Male | 1600 | 6.83 | 12 | |
| 16 | Not Cocaine Dependent | 15 | Natural | 30 | Male | 1517 | 6.37 | 11 | |
| 17 | Not Cocaine Dependent | 17 | Natural | 41 | Male | 1376 | 6 | 3 | |
| 18 | Not Cocaine Dependent | 14 | Natural | 47 | Male | 1412 | 6.49 | 3.5 | |
| 19 | Not Cocaine Dependent | 20 | Accident | 32 | Male | 1516 | 6.67 | 4 | |
| 20 | Not Cocaine Dependent | 36 | Natural | 27 | Male | 1595 | 6.55 | 3 | |
| 21 | Not Cocaine Dependent | 94 | Accident | 15 | Male | 1420 | 6.72 | 16.75 | |
| 22 | Not Cocaine Dependent | 133 | Accident | 42 | Male | 1470 | 6.75 | 2.5 | |
| 23 | Not Cocaine Dependent | 135 | Natural | 18 | Male | 1470 | 6.87 | 2 | |
| 24 | Not Cocaine Dependent | 16 | Accident | 28 | Male | 1565 | 6.32 | 2.25 | |
| 25 | Not Cocaine Dependent | 197 | Suicide | 41 | Female | 1355.2 | 6.5 | 3.5 | |
| 26 | Not Cocaine Dependent | 173 | Accident | 20 | Male | 1533 | 6.3 | 12 | |
