## Additional File 2 for "Cocaine-induced neuron subtype mitochondrial dynamics through Egr3 transcriptional regulation"

| | Forward | Reverse |
|---|---|---|
| mDrp1 | GGGCACTTAAATTGGGCTCC | TGTATTCTGTTGGCGTGGAAC |
| mNrf2 | TCTACTGAAAAGGCGGCTCA | TTGCCATCTCTGGTTTGCTG |
| mNRF1 | AGACCTCTGCTAGATTCACCG | CCTGGACTTCACAAGCACTC |
| mPolG | ACTCCTGGAACAGTTGTGCT | CGTCCATCTACTCAGGACGG |
| mTFAMSV1 | GCAGGCACTACAGCGATACA | GCCTCTACCTTTCCCATTCC |
| mTFB1 | TACGCCCTTGATAGAGCCCA | TCCTTCGAAACTGAAACGCA |
| mTom20 | CTGTGCTCTGGGCACTTAAC | AGGGTGCACACAGGTCTAAT |
| mGapdh | AGGTCGGTGTGAACGGATTTG | TGTAGACCATGTAGTTGAGGT CA |
| rDrp1 | CTCCACCTTTTGAAGCCAGG | GCAGCCGTAGTCCTCAAAGA |
| rNrf2 | CCGTACAAAACAAACACTAG CTC | AGATGGCAACGTGTTCCTTG |
| rNRF1 | ATGGCGGAAGTAATGAAAGA CG | TACTTCCCAGCAGCCTTAGC |
| rPolG | CTCCGCACCCGAAGATTTG | CCTCCTCAGAGAATGGGCAG |
| rTFAMSV1 | ATCAAGACTGTGCGTGCATC | AGAACTTCACAAACCCGCAC |
| rTFB1 | TGAAGACCCACAACTCTTTGC | CAGCAGTTAGAACCCACAGC |
| rTom20 | TGGAATGAGCCAGACACCAA | CACAGTTTGCCCTTATCCCC |
| rGapdh | TGGCCTCCAAGGAGTAAGAA | TGTGAGGGAGATGCTCAGTG |
| hDrp1 | TCTTGGAGGACTATGGCAGC | CAAAGCAGTTTGCCTGTGGA |
| hNrf2 | AGCATTGGAGTGTCAGTATGT T | ACTAGCCCAAATGGTGTCCA |
| hNRF1 | GGTGCGCTGTGGAAACAATA | CAGTAGCTCAACGCATGACC |
| hPolG | TCACCAAAGGCTCCTTGGAA | CACGGGAGCAAATACAGAGC |
| hTFAMSV1 | GGCACAGGAAACCAGTTAGG | ATGCTGGCAGAAGTCCATGA |
| hTFB1 | TGCACTACGTGGAGCTTCTT | GTCACATCTGGTCATTGGCA |
| hTom20 | TAGCCTTGTGAGCTTCGCTA | CAGCAGACGCATTCTCTCAC |
| rGapdh | TGTTCGTCATGGGTGTGAAC | GCAGGGATGATGTTCTGGAG |
