## Additional File 3 for "Cocaine-induced neuron subtype mitochondrial dynamics through Egr3 transcriptional regulation"

| | Species | Nrf1 | Nrf2 | Poly | Tfam | Tfb1 | Tomm20 | Egr3 | PGC1$\alpha$ | Drp1 |
|---|---|---|---|---|---|---|---|---|---|---|
| **Total Brain** | human | - | ↑ | - | - | - | - | N/A | ↑ | ↑[3] |
| | rat (self-admin) | ↑ | - | - | - | ↑ | ↑ | ↓[1] | - | ↑[3] |
| | mouse (i.p.) | ↑ | ↑ | - | - | - | ↑ | ↓[1] | ↑[2] | ↑[3] |
| **D1 MSN** | mouse (i.p.) | - | - | - | ↑ | - | - | ↑[1] | ↑[2] | ↑[3] |
| **D2 MSN** | mouse (i.p.) | - | - | ↓ | ↓ | ↓ | ↓ | ↓[1] | ↓[2] | ↓[3] |

[1]Chandra, R., et al., (2015). *Journal of Neuroscienc*e, 35(20), 7927–7937
[2]Chandra, R., et al., (2017). *Biological Psychiatry*, 81(7), 564-572
[3]Chandra, R., et al., (2017). Neuron*, 96(6), 1327-1341.e6*
