## Additional File 4 for "Cocaine-induced neuron subtype mitochondrial dynamics through Egr3 transcriptional regulation"

| Figure | Test | Test Statistic | N Per Group and Effect Sizes |
|---|---|---|---|
| 1a Drp1 | Unpaired T-test | $t_{(10)}$=2.3,P=0.0448 | 6 Saline, 6 Cocaine; Cohens d=1.3238 |
| 1a Nrf1 | Unpaired T-test | $t_{(12)}$=1.01,P=0.3335 | 7 Saline, 7 Cocaine; Cohens d=0.538 |
| 1a Nrf2 | Unpaired T-test | $t_{(10)}$=2.27,P=0.0467 | 5 Saline, 7 Cocaine; Cohens d=1.001 |
| 1a Polγ | Unpaired T-test | $t_{(11)}$=2.33,P=0.0406 | 6 Saline, 7 Cocaine; Cohens d=0.992 |
| 1a Tfam | Unpaired T-test | $t_{(3)}$=1.13,P=0.3423 | 3 Saline, 4 Cocaine; Cohens d=1.162 |
| 1a Tfb1 | Unpaired T-test | $t_{(4)}$=0.83,P=0.4548 | 3 Saline, 3 Cocaine; Cohens d=0.675 |
| 1a Tomm20 | Unpaired T-test | $t_{(10)}$=1.7,P=0.1214 | 5 Saline, 7 Cocaine; Cohens d=0.830 |
| 2a Nrf1 | Unpaired T-test | $t_{(11)}$=3.37,P=0.0064 | 6 Saline, 7 Cocaine; Cohens d=1.940 |
| 2a Nrf2 | Unpaired T-test | $t_{(10)}$=4.67,P=0.0009 | 6 Saline, 6 Cocaine Cohens d=2.6963 |
| 2a Polγ | Unpaired T-test | $t_{(10)}$=1.15,P=0.2797 | 6 Saline, 6 Cocaine; Cohens d=0.660 |
| 2a Tfam | Unpaired T-test | $t_{(11)}$=1.83,P=0.0952 | 6 Saline, 7 Cocaine; Cohens d=1.052 |
| 2a Tfb1 | Unpaired T-test | $t_{(10)}$=1.6,P=0.1414 | 6 Saline, 6 Cocaine; Cohens d=0.922 |
| 2a Tomm20 | Unpaired T-test | $t_{(10)}$=2.76,P=0.0203 | 5 Saline, 7 Cocaine; Cohens d=1.705 |
| 2b Nrf1 | Unpaired T-test | $t_{(9)}$=2.71,P=0.0244 | 5 Saline, 6 Cocaine; Cohens d=1.606 |
| 2b Nrf2 | Unpaired T-test | $t_{(9)}$=1.92,P=0.0884 | 5 Saline, 6 Cocaine; Cohens d=1.168 |
| 2b Pgc1α | Unpaired T-test | $t_{(9)}$=1.93,P=0.0866 | 5 Saline, 6 Cocaine; Cohens d=1.202 |
| 2b Polγ | Unpaired T-test | $t_{(9)}$=1.86,P=0.0961 | 5 Saline, 6 Cocaine; Cohens d=1.106 |
| 2b Tfam | Unpaired T-test | $t_{(9)}$=0.66,P=0.5292 | 5 Saline, 6 Cocaine; Cohens d=0.383 |
| 2b Tfb1 | Unpaired T-test | $t_{(9)}$=2.44,P=0.0379 | 5 Saline, 6 Cocaine; Cohens d=1.441 |
| 2b Tomm20 | Unpaired T-test | $t_{(9)}$=2.49,P=0.0348 | 5 Saline, 6 Cocaine; Cohens d=1.475 |
| 2c Tfb1 | Unpaired T-test | $t_{(27)}$=1.45,P=0.1599 | 14 Control, 15 Cocaine dependents; Cohens d=0.528 |
| 2c Nrf1 | Unpaired T-test | $t_{(28)}$=1.43,P=0.1649 | 15 Control, 15 Cocaine dependents; Cohens d=0.521 |
| 2c Pgc1α | Unpaired T-test | $t_{(28)}$=2.05,P=0.0505 | 15 Control, 15 Cocaine dependents; Cohens d=0.746 |
| 2c Nrf2 | Unpaired T-test | $t_{(23)}$=2.33,P=0.0296 | 12 Control, 13 Cocaine dependents; Cohens d=0.937 |
| 2c Polγ | Unpaired T-test | $t_{(26)}$=2.12,P=0.044 | 13 Control, 15 Cocaine dependents; Cohens d=0.779 |
| 2c Tfam | Unpaired T-test | $t_{(29)}$=0.29,P=0.7785 | 16 Control, 15 Cocaine dependents; Cohens d=0.1024 |
| 2c Tomm20 | Unpaired T-test | $t_{(28)}$=1.27,P=0.2149 | 15 Control, 15 Cocaine dependents; Cohens d=0.463 |
| 3b Nrf1 | Unpaired T-test | $t_{(8)}$=1.14,P=0.2883 | 4 Saline, 6 Cocaine; Cohens d=0.810 |
| 3b Nrf2 | Unpaired T-test | $t_{(9)}$=2.05,P=0.071 | 5 Saline, 6 Cocaine; Cohens d=1.272 |
| 3b Polg | Unpaired T-test | $t_{(7)}$=0.66,P=0.5358 | 4 Saline, 5 Cocaine; Cohens d=0.448 |
| 3b TFAM | Unpaired T-test | $t_{(6)}$=3.38,P=0.015 | 4 Saline, 4 Cocaine; Cohens d=2.388 |
| 3b TFB1 | Unpaired T-test | $t_{(10)}$=1.61,P=0.1402 | 6 Saline, 6 Cocaine; Cohens d=0.925 |
| 3b Tomm20 | Unpaired T-test | $t_{(9)}$=2.24,P=0.0526 | 6 Saline, 5 Cocaine; Cohens d=1.318 |
| 3c Nrf1 | Unpaired T-test | $t_{(9)}$=1.95,P=0.0836 | 5 Saline, 6 Cocaine; Cohens d=1.182 |
| 3c Nrf2 | Unpaired T-test | $t_{(9)}$=2.01,P=0.0762 | 5 Saline, 6 Cocaine; Cohens d=1.181 |

| | | | |
|---|---|---|---|
| 3c Polg | Unpaired T-test | $t_{(9)}$=2.47,P=0.0362 | 5 Saline, 6 Cocaine; Cohens d=1.523 |
| 3c TFAM | Unpaired T-test | $t_{(9)}$=3.34,P=0.0088 | 5 Saline, 6 Cocaine; Cohens d=2.050 |
| 3c TFB1 | Unpaired T-test | $t_{(9)}$=2.32,P=0.0459 | 5 Saline, 6 Cocaine; Cohens d=1.431 |
| 3c Tomm20 | Unpaired T-test | $t_{(8)}$=5.41,P=0.0007 | 5 Saline, 5 Cocaine; Cohens d=3.421 |
| 4b Pgc1α | Two-way ANOVA | Drug: $F_{(1,18)}$=4.869, P=0.0406, Virus: $F_{(1,18)}$=4.257, P=0.0538, Interaction: $F_{(1,18)}$ = 9.398, p = 0.0067. | 5 SS-miR Saline, 6 SS-miR Cocaine, 6 Egr3-miR Saline, 6 Egr3-miR Cocaine; Cohens d=0.772 |
| 4b Drp1 | Two-way ANOVA | Drug: $F_{(1,20)}$=4.511, P=0.0463, Virus: $F_{(1,20)}$=12.67, P=0.0020, Interaction: $F_{(1,20)}$=12.67, p=0.0020, | 5 SSmiR-Saline, SSmiR-Cocaine, 6 Egr3miR-Saline, 7 Egr3miR; Cohens d=1.214 |
| 4d frequency | Two-way ANOVA | Drug: $F_{(5,70)}$=73.19, P<0.0001, Virus: $F_{(3,14)}$=0.6858, P=0.5754, Interaction: $F_{(15, 70)}$=3.035, P=0.0009 | 4 SS-miR Saline , 4 SS-miR Cocaine 5 Egr3-miR Saline, 5 Egr3-miR Cocaine, |
| 4d density | Two-way ANOVA | Drug: $F_{(1,14)}$= 3.441, P=0.0848, Virus: $F_{(1,14)}$=2.207, P=0.1595, Interaction: $F_{(1,14)}$=0.2823, P=0.6035 | 4 SS-miR Saline , 4 SS-miR Cocaine 5 Egr3-miR Saline, 5 Egr3-miR Cocaine, |
| 4d index | Two-way ANOVA | Drug $F_{(1,14)}$=0.01078, P = 0.9188, Virus $F_{(1,14)}$=0.3075, P=0.5880, Interaction: $F_{(1,14)}$=3.919, P= 0.0678. | 4 SS-miR Saline , 4 SS-miR Cocaine 5 Egr3-miR Saline, 5 Egr3-miR Cocaine, |
| 4d volume ratio | Two-way ANOVA | Drug: $F_{(1,14)}$=3.008, P=0.1048, Virus: $F_{(1,14)}$=0.08972, P=0.7689, Interaction: $F_{(1,14)}$=3.919, P=0.0678. | 4 SS-miR Saline , 4 SS-miR Cocaine 5 Egr3-miR Saline, 5 Egr3-miR Cocaine, |
